# The CWI pathway is activated through high hydrostatic pressure, enhancing glycerol efflux via the aquaglyceroporin Fps1 in *Saccharomyces cerevisiae*

**DOI:** 10.1101/2022.11.15.516693

**Authors:** Takahiro Mochizuki, Toshiki Tanigawa, Seiya Shindo, Momoka Suematsu, Yuki Oguchi, Tetsuo Mioka, Yusuke Kato, Mina Fujiyama, Eri Hatano, Masashi Yamaguchi, Hiroji Chibana, Fumiyoshi Abe

## Abstract

The fungal cell wall is the initial barrier for the fungi against diverse external stresses, such as osmolarity changes, harmful drugs, and mechanical injuries. This study explores the roles of osmoregulation and the cell wall integrity (CWI) pathway in response to high hydrostatic pressure in the yeast *Saccharomyces cerevisiae*. We demonstrate the roles of the transmembrane mechanosensor Wsc1 and aquaglyceroporin Fps1 in a general mechanism to maintain cell growth under high-pressure regimes. The promotion of water influx into cells at 25 MPa, as evident by an increase in cell volume and a loss of the plasma membrane eisosome structure, promotes the activation of Wsc1, an activator of the CWI pathway. The downstream mitogen-activated protein kinase Slt2 was hyperphosphorylated at 25 MPa. Glycerol efflux increases via Fps1 phosphorylation, which is initiated by downstream components of the CWI pathway and contributes to the reduction in intracellular osmolarity under high pressure. The elucidation of the mechanisms underlying adaption to high pressure through the well-established CWI pathway could potentially translate to mammalian cells and provide novel insights into cellular mechanosensation.

## Introduction

Microorganisms encounter a wide variety of extreme environmental conditions, including heat, cold, salinity, or nutrient starvation. These stresses are first detected by sensor proteins that trigger the expression of specific genes via intracellular signaling pathways to respond to and overcome a crisis. Eukaryotes are also subjected to physical stresses, including tension, shear forces, and hydrostatic pressure. In mammalian cells, forces are detected, at the cell surface, through specific classes of integral membrane receptors, such as integrins and cadherins, which transduce stimuli into biochemical signals that initiate adaptive responses (Kechagia *et al.*, 2019; Ladoux and Mege, 2017). Unlike stretching or expansion, hydrostatic pressure acts on cells as a uniform stress and affects all substances and reactions in the cell, according to Pascal’s principle. Therefore, it is difficult to identify sensors of high hydrostatic pressure and determine their effects on complex intracellular signaling networks. Nevertheless, there is ample evidence that cells modify their metabolism, transcription, translation, or post-translational processes in response to increasing hydrostatic pressure (Abe *et al.*, 1999; Abe, 2021; Bourges *et al.*, 2017; Hodder *et al.*, 2020; Iwahashi, 2015; Mishra *et al.*, 2022; Pattappa *et al.*, 2019).

Human cells also experience high hydrostatic pressures. In joints, cartilage is typically subjected to hydrostatic pressures between 3–10 MPa (0.1 MPa = 1 bar = 0.9869 atm = 1.0197 kg/cm^2^) (Afoke *et al.*, 1987), with a maximum pressure of 18 MPa in the hip joint (Hodge *et al.*, 1989). The loading of endothelial cells with a low pressure of 50 mmHg (~6.7 kPa) activates protein kinase C and transiently activates the Ras/ERK pathway as a result of the water efflux through aquaporin 1 (Yoshino *et al.*, 2020). High-pressure adaptation mechanisms in deep-sea microorganisms have also received considerable attention. However, little is known regarding how microbial cells detect such stimuli and convert them to intracellular signals.

Although *S. cerevisiae* is not a deep-sea piezophile, it can be used to elucidate the molecular mechanisms underlying the cell’s responses to high hydrostatic pressure by applying basic knowledge of genetics and protein functions (Abe, 2021). Pressure pretreatment at a sublethal level (50 MPa for 1 h) increased the viability of yeast cells at 200 MPa, which is governed by two stress-induced transcription factors, Msn2 and Msn4 (Domitrovic *et al.*, 2006). High pressure of 25 MPa accumulates superoxide anions in mitochondria in a mutant deficient in superoxide dismutase 1 (Sod1), and the mutant shows marked sensitivity to high pressures (Funada *et al.*, 2022). Incubation of yeast cells at 25 MPa for 5 h resulted in the upregulation of the *DAN/TIR* family mannoprotein genes, which are induced under hypoxic conditions and at low temperatures (Abe, 2007). Target of rapamycin complex 1 (TORC1), an evolutionarily conserved serine/threonine kinase in eukaryotes, was activated by a pressure of 25 MPa (Uemura *et al.*, 2020). We demonstrated that TORC1 plays a critical role in maintaining an appropriate glutamine level under high pressure by downregulating the *de novo* synthesis of glutamine (Uemura *et al.*, 2020). Such evidence indicates that high-pressure stress induces diverse changes in intracellular processes and that cells survive by adaptively coping with each change. It may appear that yeasts are not exposed to high hydrostatic pressure (equivalent to tens of MPa) on earth, except for those inhabiting the deep sea. However, microorganisms inhabiting soil that are trampled by a running 4 ton African elephant also experience comparable pressure. In addition, giant dinosaurs weighing up to 100 tons roamed on the earth hundreds of millions of years ago, hence it must have been crucial for soil microorganisms to acquire tolerance to extremely high pressure to survive during evolution.

The fungal cell wall provides the first barrier against external stresses, such as osmolarity changes, harmful drugs, and mechanical injuries. It is an elastic structure with a thickness of 50–500 nm and is composed of β1,3-glucan, β1,6-glucan, chitin, and mannoproteins (Lesage and Bussey, 2006; Levin, 2011; Orlean, 2012). The Young’s modulus of the cell wall is estimated to be between 10 and 100 MPa (Davi *et al.*, 2019; Ma *et al.*, 2005), which is an intermediate value between that of low-density polyethylene and rubber. The primary mechanical role of the cell wall is to maintain cell morphology by balancing the tension generated by the turgor pressure, which is important for polarized cell growth (Bi and Park, 2012; Madden and Snyder, 1998; Pruyne and Bretscher, 2000). In *S. cerevisiae,* cell wall destabilization is sensed by transmembrane sensors of the WSC family (Wsc1, Wsc2, and Wsc3), as well as Mid2 and its homolog Mtl1 (Jimenez-Gutierrez *et al.*, 2020; Kock *et al.*, 2015; Levin, 2005; Levin, 2011; Philip and Levin, 2001), which activates a conserved signal transduction cascade, namely, the cell wall integrity (CWI) pathway. Their architecture is analogous to that of the mammalian transmembrane integrins (Elhasi and Blomberg, 2019). Among the five sensor proteins in *S. cerevisiae*, Wsc1 has been extensively studied both genetically and biophysically (Banavar *et al.*, 2018; Dupres *et al.*, 2009; Heinisch *et al.*, 2010; Merchan *et al.*, 2004; Neeli-Venkata *et al.*, 2021; Philip and Levin, 2001; Schoppner *et al.*, 2022). The CWI pathway in *S. cerevisiae* is activated in response to cell wall damage; however, recent evidence suggests that it is also involved in other stress conditions, indicating functional diversity (Banavar *et al.*, 2018; Cruz *et al.*, 2013; Kock *et al.*, 2016; Straede and Heinisch, 2007). To date, the effects of high hydrostatic pressure in yeast cells have not been studied with a focus on osmoregulation and the CWI pathway.

Atomic force microscopy (AFM) analysis revealed that a serine-threonine-rich (STR) domain has a nanospring-like structure that expands and contracts under external pressure, which in turn causes dephosphorylation of the cytoplasmic tail of Wsc1, transmitting signals downstream (Dupres *et al.*, 2009; Heinisch *et al.*, 2010; Vay *et al.*, 2004). The cytoplasmic tail of Wsc1 interacts with the GDP/GTP exchange factor Rom2, which subsequently activates small GTPase Rho1(Philip and Levin, 2001; Vay *et al.*, 2004). Bem2, Sac7, Lrg1, and Bag7 act as GTPase-activating proteins (GAPs) for Rho1 (Fig. 1)(Martin *et al.*, 2000; Peterson *et al.*, 1994; Schmidt *et al.*, 1997; Schmidt *et al.*, 2002). Rho1 regulates protein kinase C (Pkc1), a serine/threonine protein kinase that is essential for cell wall remodeling (Levin, 2011).

**Fig. 1.**
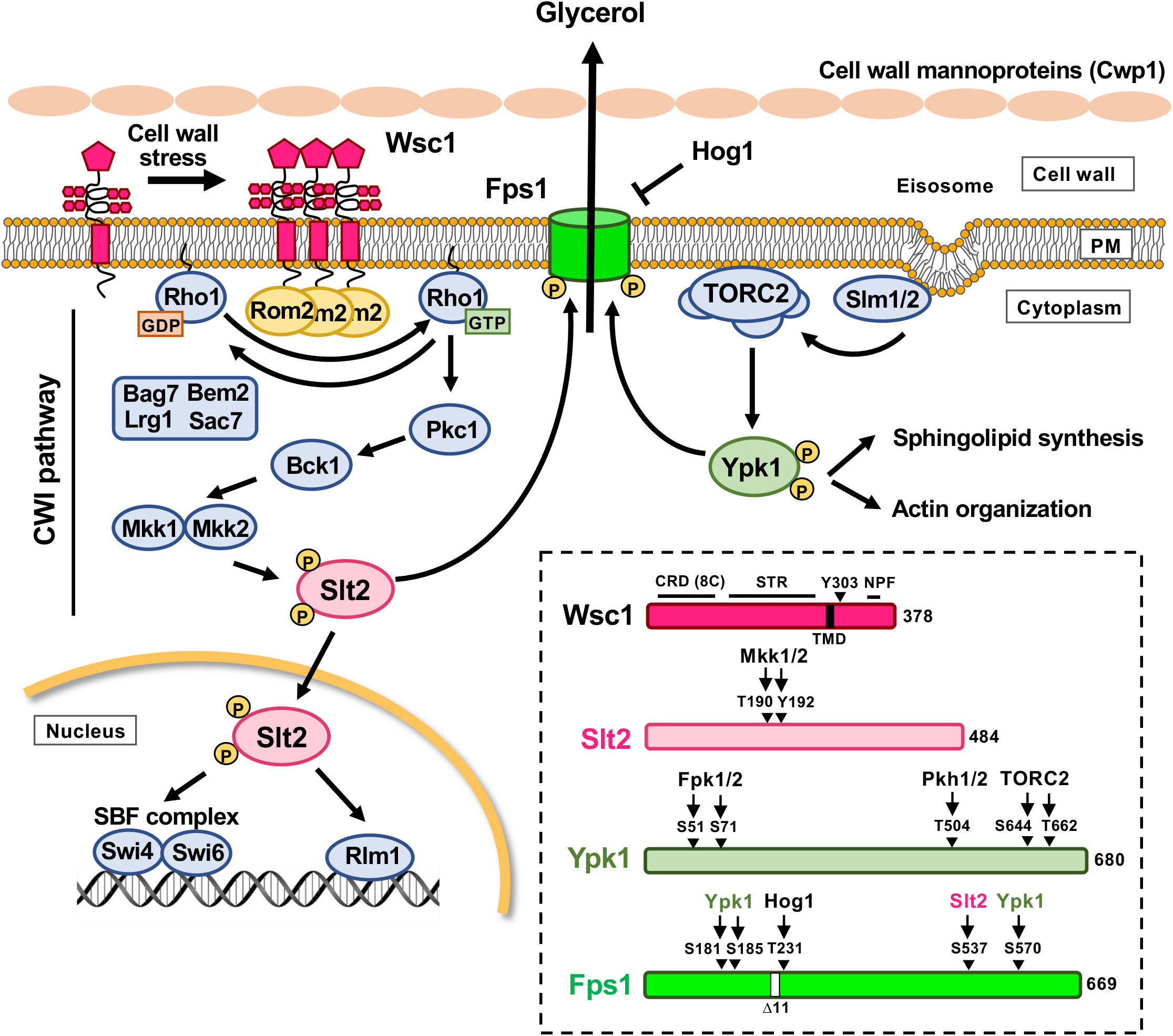
Schematic representation of the CWI pathway in *Saccharomyces cerevisiae*. Five cell wall sensor proteins are depicted. Upon cell wall stress, Wsc1 first clusters at the cell surface. Binding of the GDP/GTP exchanging factor, Rom2, to the cytoplasmic tail of Wsc1 activates the small GTPase, Rho1. Rho1-GTP activates the protein kinase C, Pkc1, which activates a conserved mitogen-activated protein kinase (MAPK) cascade. The cascade consists of Bck1 (MAPKKK), Mkk1/Mkkk2 (MAPKKs), and Slt2 (MAPK). Phosphorylated Slt2 translocates into the nucleus and promotes cell cycle progression through the SBF complex and transcription of genes encoding cell wall proteins by Rlm1. Slt2 also phosphorylates the aquaglyceroporin, Fps1, and promotes Fps1 opening to expel glycerol. When cells are stressed, tension is applied to the plasma membrane, causing the loss of the eisosome, a plasma membrane invaginating structure. Slm1/Slm2 localized in the eisosome migrate to TORC2. Consequently, TORC2 is activated and phosphorylates Ypk1. Phosphorylated Ypk1 also phosphorylates Fps1, positively regulating glycerol efflux. The phosphorylation sites, kinases, and functional motifs in major proteins used in this study are outlined below. CRD (8C), cysteine-rich domain (C27, C49, C53, C69, C71, C86, C90, and C98); STR, serine/threonine-rich domain (111–265 amino acid residues); NPF motif, N344-P345-F346.

Pkc1 then activates three protein kinase-signaling modules that integrate the mitogen-activated protein kinase kinase kinase (MAPKKK) Bck1, redundant mitogen-activated protein kinase kinases (MAPKKs) Mkk1 and Mkk2, and mitogen activated protein kinase (MAPK) Slt2 (Levin, 2011). Slt2 kinase regulates many cellular processes, such as fine-tuning of the CWI pathway, transcriptional response to cell wall damage, cell cycle progression, and nuclear export of mRNA (Gonzalez-Rubio *et al.*, 2022), and mediates aquaglyceroporin Fps1 phosphorylation (Ahmadpour *et al.*, 2016; Levin, 2005; Mollapour *et al.*, 2009). Although the CWI cascade is well characterized (Fig. 1), the exact mode of Wsc1 action in response to different environmental cues remains unclear.

Fps1 is under the control of the CWI pathway (Ahmadpour *et al.*, 2016; Laz *et al.*, 2020). In yeast, when the cell wall is damaged, water flows into the cell due to the osmotic pressure difference between the inside and outside of the cell. To cope with this stress and prevent osmolysis, yeast cells activate Fps1 to expel glycerol and eliminate osmotic pressure differences (Tamas *et al.*, 1999). The opening and closing of Fps1 is a complex process mediated by protein phosphorylation involving several signaling pathways (Fig. 1). First, MAP kinase Hog1 is activated in response to hyperosmotic stress and phosphorylates the T231 residue of Fps1, resulting in Fps1 closure (Mollapour and Piper, 2007; Thorsen *et al.*, 2006). Hog1 negatively regulates this channel by displacing the Fps1-activating proteins Rgc1 and Rgc2, thereby causing Fps1 closure (Lee *et al.*, 2013). Second, the open state of Fps1 requires phosphorylation by Ypk1, a target of TORC2-dependent protein kinase (Muir *et al.*, 2015). Third, Fps1 and Slt2 physically interact via phosphorylation of the S537 residue of Fps1, which promotes the opening of the glycerol channel (Ahmadpour *et al.*, 2016).

Remarkably, while imaging wild-type *S. cerevisiae* cells, minor cell swelling was observed after high-pressure incubation at 25 MPa. This suggests that high pressure may facilitate the influx of water into the cell or damage the cell wall, further allowing water influx. This observation steered us to study high-pressure signaling, with a focus on water influx, turgor pressure, the CWI pathway via Wsc1, MAP kinase Slt2, and aquaglyceroporin Fps1 (Fig. 1). Our study provides novel insights into the signaling events triggered by the application of high-pressure and the resulting molecular mechanisms that help cells to cope with such stress.

## Results

### Wsc1 is required for efficient cell growth under high pressure

Based on the previous studies (Jimenez-Gutierrez *et al.*, 2020; Kock *et al.*, 2015; Levin, 2005; Levin, 2011; Philip and Levin, 2001), we hypothesized that any of the five sensor proteins, Wsc1–3, Mid2, and Mtl1, could detect hydrostatic pressure changes, either directly or as a result of cell wall damage, and activate the CWI pathway to allow for high-pressure growth. Cells were cultured in synthetic complete (SC) medium at an OD_600_ of 0.01 for 48 h, under atmospheric pressure (0.1 MPa) or 25 MPa. We found that the *wsc1*Δ mutant exhibited a moderate growth defect at 25 MPa, whereas the other mutants grew as efficiently as the wild-type cells (Fig. 2A). The result suggests that Wsc1 plays a primary role in high-pressure growth, while the other sensor proteins may contribute to the growth with overlapping functions.

**Fig. 2.**
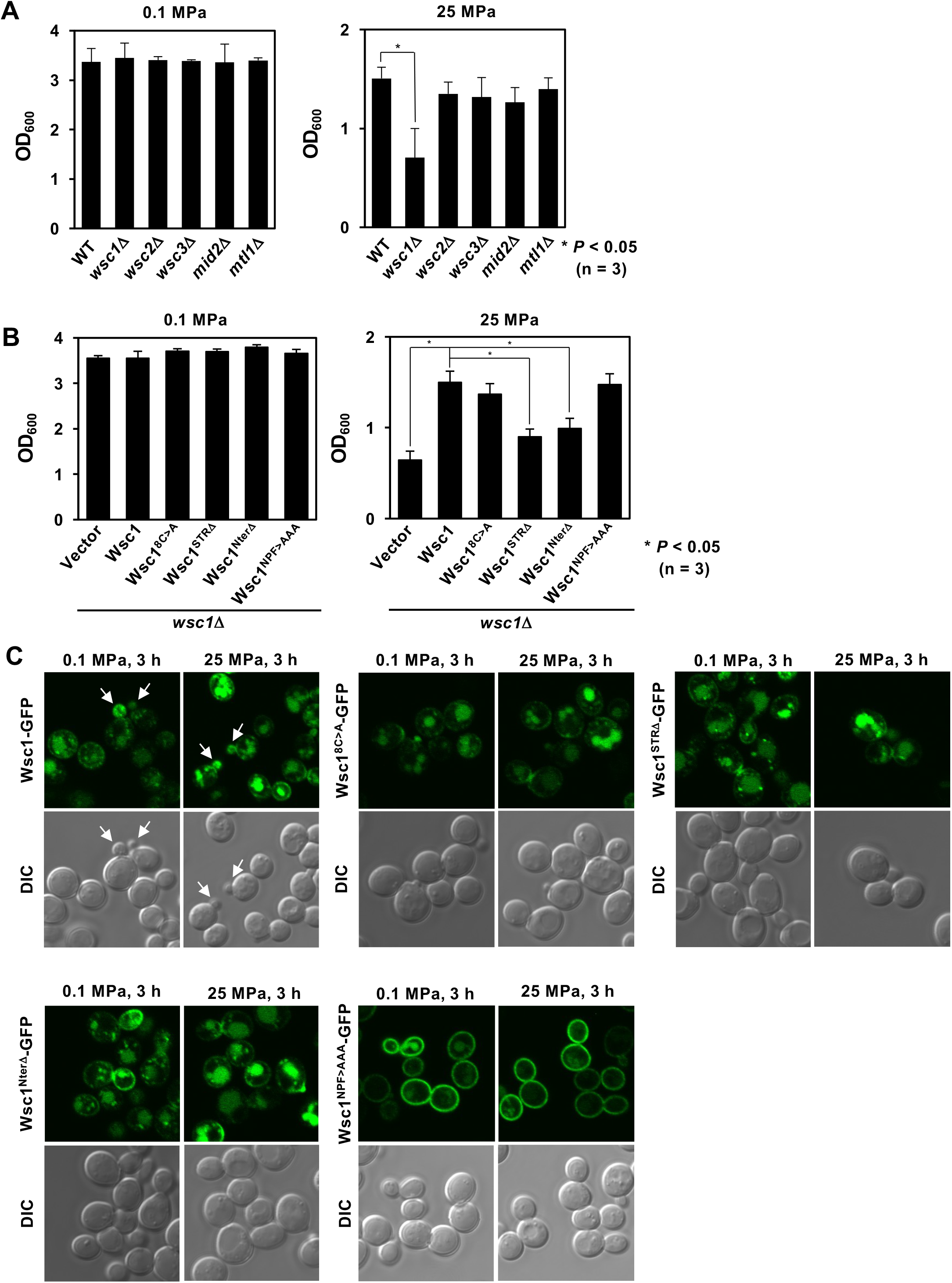
Wsc1 is required for growth under high pressure. (A) Wild-type and mutant cells were cultured in SC medium at 0.1 MPa or 25 MPa for 48 h with an initial OD600 = 0.01. Unless otherwise specified, the high-pressure growth test was carried under this condition throughout the experiments. (B) Cells with mutations in the extracellular domain of Wsc1 or the NPF>AAA substitution were cultured in SC medium at 0.1 or 25 MPa. (C). Wild-type cells expressing Wsc1-GFP or the mutant forms were cultured at 0.1 or 25 MPa for 3 h and were observed under a confocal laser microscope immediately after depressurization. Data are presented as mean ± standard deviation of three independent experiments.

Wsc1 is characterized by an N-terminal extracellular cysteine-rich domain (CRD), followed by a glycosylated STR domain, a single transmembrane domain, and a C-terminal cytoplasmic tail, which is considered to be the mechanosensory region (Fig. 1). The CRD and STR domains detect structural changes in the cell wall (Dupres *et al.*, 2009; Heinisch *et al.*, 2010). An alanine substitution of eight cysteines in the CRD of Wsc1 had a marginal effect on high-pressure growth ability, whereas deletions of the STR and the entire N-terminal region caused sensitivity to high pressure (Fig. 2B). These results suggest that the STR of Wsc1 can detect structural changes in the cell wall caused by high pressure.

Single-molecule AFM results indicated that Wsc1 assembles in patches as large as 200 nm on the plasma membrane, thereby enhancing transduction of the CWI signaling pathway (Dupres *et al.*, 2009; Heinisch *et al.*, 2010). Wsc1 assembles at the tips and bud necks where cells undergo polar growth, and endocytosis is essential for their polarity (Piao *et al.*, 2007). To determine the effect of high pressure on the distribution of Wsc1, we observed the localization of Wsc1-GFP before and after culturing the cells at 25 MPa. Cells were observed after depressurization. As previously reported (Piao *et al.*, 2007; Wilk *et al.*, 2010), Wsc1-GFP localized mainly to bud tips, but also to the plasma membrane of mother cells as patchy structures, and possibly to late endosomes as punctate structures and vacuoles (Fig. 2C). The bud tip localization of Wsc1-GFP was maintained after the high-pressure incubation although its vacuolar degradation was slightly enhanced (Fig. 2C). Enhanced vacuolar degradation of membrane proteins under high pressure has also been observed in the tryptophan permeases Tat1 and Tat2 (Abe and Iida, 2003; Suzuki *et al.*, 2013). The overall localization of GFP-tagged Wsc1^8C>A^, Wsc1^STRΔ^, and Wsc1^NterΔ^ was comparable with the wild-type Wsc1-GFP (Fig. 2D). The polarized distribution of Wsc1 to bud tips requires endocytosis (Piao *et al.*, 2007; Wilk *et al.*, 2010). The Wsc1 mutant, in which the NPF sequence at the C-terminal tail was replaced by AAA, exhibited a uniform distribution of Wsc1 within the plasma membrane (Piao *et al.*, 2007). Wsc1^NPF>AAA^-GFP was present throughout the plasma membrane (Fig. 2C) and Wsc1^NPF>AAA^ expressing cells grew as efficiently as the wild-type cells at 25 MPa (Fig. 2B). Therefore, we suggest that Wsc1 is required for efficient cell growth under high pressure but does not need to be exclusively localized to the bud tips.

### Wsc1 activates the CWI pathway in response to high pressure

To address the role of Wsc1 in optimal growth under high pressure, we focused on the downstream effector Rom2. The Wsc1 C-terminal cytoplasmic domain residue, Y303, is critical for its interaction with Rom2 (Vay *et al.*, 2004). Both the disruption of *ROM2* and the Wsc1-Y303A mutation, which interferes with the Wsc1–Rom2 interaction, resulted in growth defects at 25 MPa (Fig. 3A). Among the four Rho1 GAP genes, deletions in *BEM2, SAC7*, and *LRG1* caused growth defects at 25 MPa to varying degrees (Fig. 3A). The expression of GTP-bound active forms of Rho1-F35L or Q68H failed to restore the high-pressure sensitivity of the *rom2*Δ mutant (Fig. 3B), suggesting that the association of Rho1 with Rom2 is critically important for high-pressure growth. The expression of the constitutively active form of Pkc1-R398P allowed the *rom2*Δ mutant to grow efficiently at 25 MPa (Fig. 3B). Disruption of *PKC1*, *BCK1*, or *SLT2* resulted in severe high-pressure sensitivity, whereas only disruption of *MKK1*, and not *MKK2*, caused moderate high-pressure sensitivity (Fig. 3C). These results suggest that Wsc1 acts as a mechanosensor that detects hydrostatic pressure changes and may transmit the pressure signal into the CWI pathway.

**Fig. 3.**
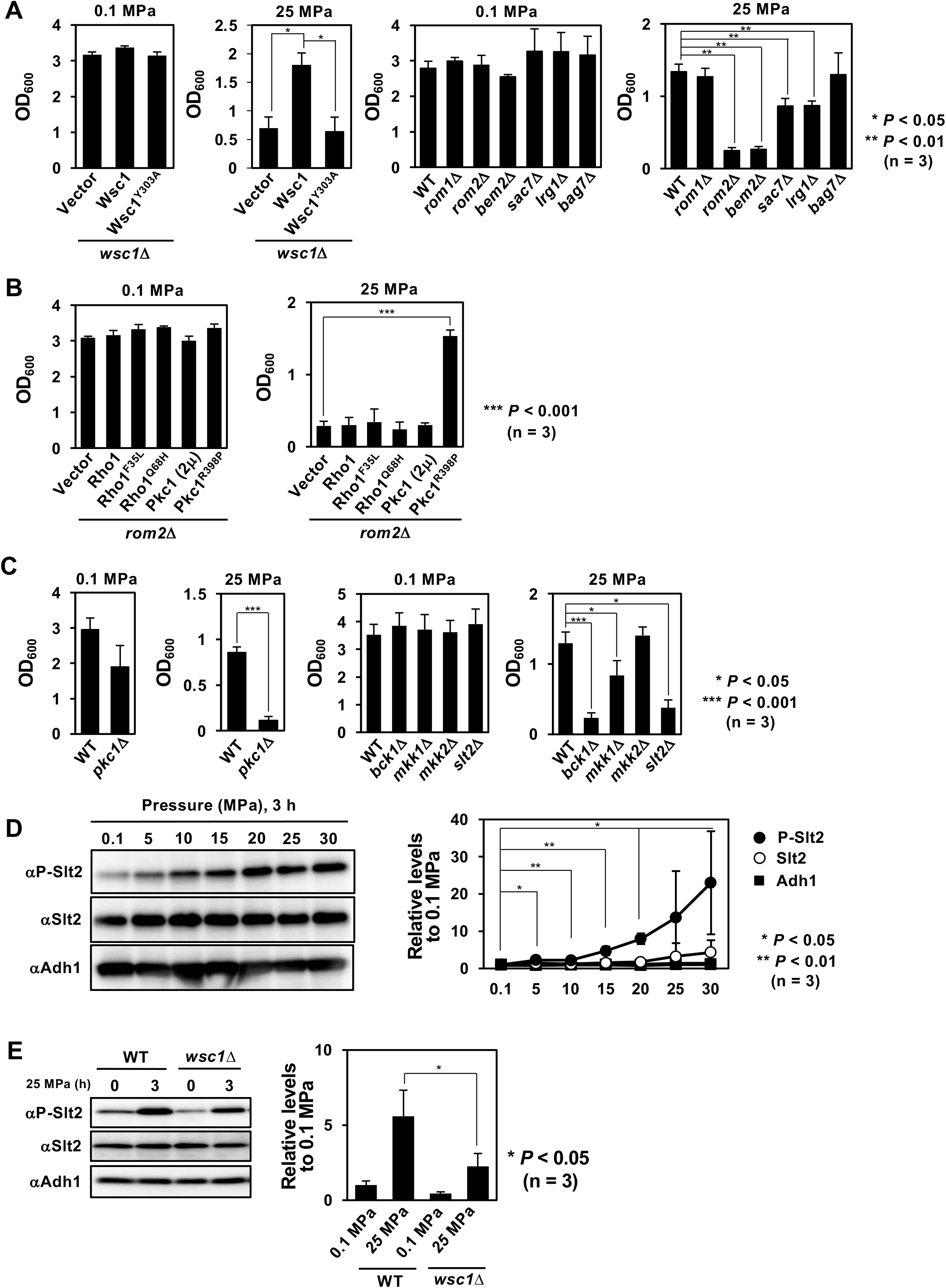
CWI pathway is activated by high pressure. (A) Mutant strains with various defects in the CWI pathway were cultured in SC medium at 0.1 or 25 MPa. (B) *rom2*Δ cells expressing Rho1 or Pck1 derivatives were cultured in SC medium at 0.1 or 25 MPa. (C) Mutant strains with various defects in the CWI pathway were cultured in SC medium at 0.1 or 25 MPa. Strains DL100 and DL376 harboring YCplac22 (*TRP1* and *CEN*) were used to examine the *PKC1* deficiency (left). (D) Wild-type cells were cultured in SC medium under various pressure conditions for 3 h. Slt2 and phosphorylated Slt2 (P-Slt2) were detected using western blotting using specific antibodies. The relative intensities, P-Slt2/Adh1 and Slt2/Adh1, were quantified in an ImageQuant LAS4000 mini. (E) Wild-type or *wsc1*Δ cells were cultured in SC medium at 0.1 or 25 MPa for 3 h. Western blotting was performed as in (D). Data are presented as mean ± standard deviation of three independent experiments.

To validate this hypothesis, we assessed the phosphorylation of Slt2 in response to high pressure and found it to increase in a pressure-dependent manner, suggesting that high pressure activates the CWI pathway (Fig. 3D). Also, disruption of *WSC1* reduced the phosphorylation level of Slt2 by 60% at 25 MPa compared with the wild-type level. This indicates that other proteins, such as Mid2, may also activate the CWI pathway in response to pressure (Fig. 3E). In contrast, loss of *BCK1* resulted in a complete loss of Slt2 phosphorylation (data not shown).

Slt2 is under the control of the CWI signaling pathway, which is responsible for maintaining cell wall homeostasis (Gonzalez-Rubio *et al.*, 2022; Levin, 2005; Martin *et al.*, 2000). Thus, hyperphosphorylated Slt2 may modulate the transcription of downstream genes through phosphorylating its substrates under high pressures. To verify this, we analyzed the effect of high pressure on the localization of Swi4, one of the downstream transcription factors of Slt2. Wild-type cells that express Swi4-GFP were incubated with 15 µg/mL nocodazole in a YPD medium for 2 h at 0.1 MPa to induce cell cycle arrest in the G_2_/M phase (Kim *et al.*, 2010) when most of the Swi4-GFP was present in the cytoplasm. Synthetic Complete (SC) medium was not used in this analysis because nocodazole was ineffective in arresting the cell cycle even at higher concentrations (data not shown). Cells were then cultured for 5 h at 0.1 MPa or 25 MPa. We found that the percentage of cells with Swi4-GFP migrating from the cytoplasm to the nucleus was higher at 25 MPa (46%, n > 300) than at 0.1 MPa (17%, n > 300) (Fig. 4A). Furthermore, cells lacking Swi4 or Swi6 showed marked high-pressure sensitivity (Fig. 4B). These results support our model, which confirms that high pressure activates the CWI pathway.

**Fig. 4.**
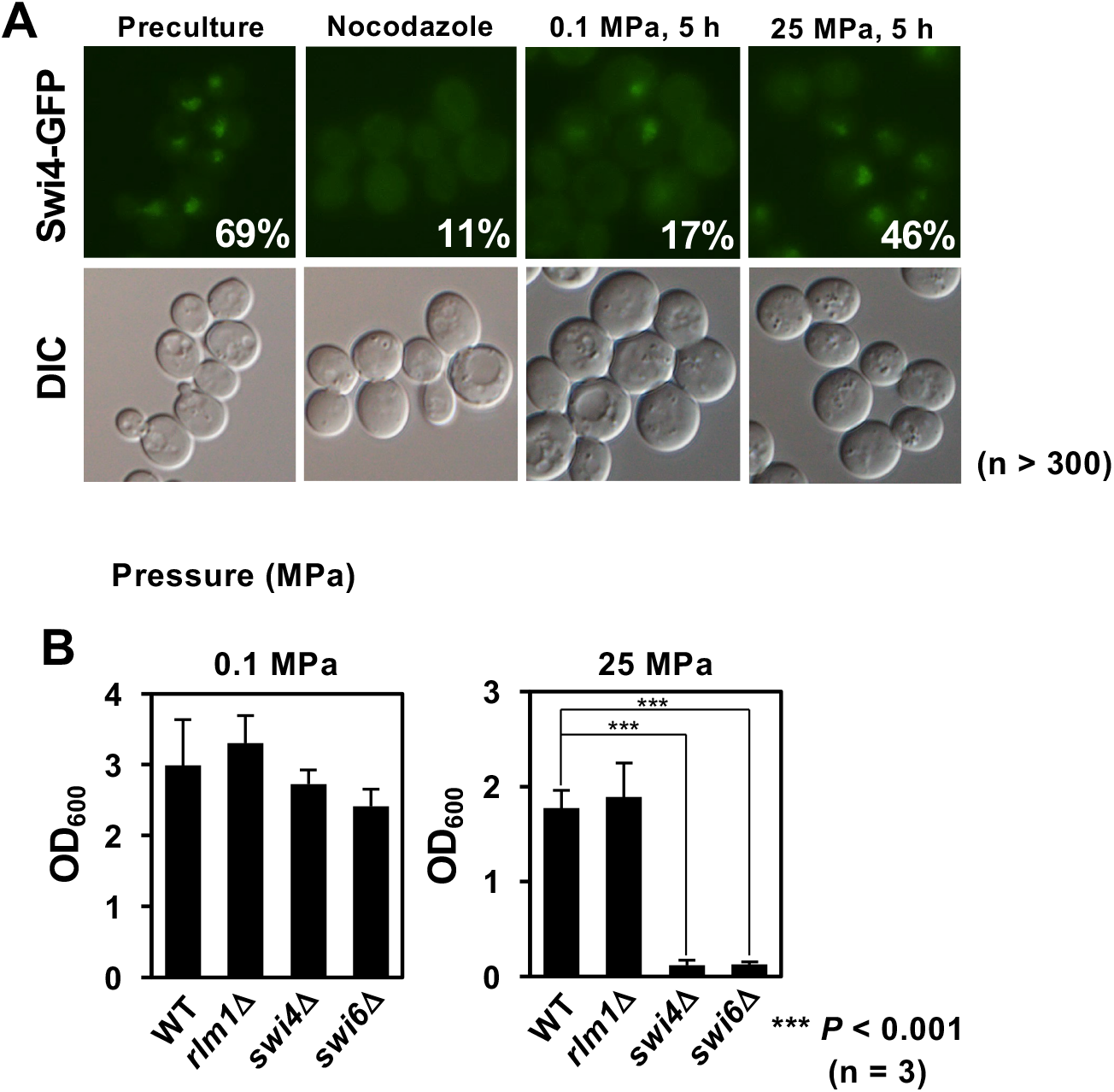
High pressure promotes nuclear translocation of Swi4. (A) Wild-type cells expressing Swi4-GFP were incubated with 15 µg/mL nocodazole for 2 h at 0.1 MPa. The cells were then cultured for another 5 h at 0.1 or 25 MPa and were observed after depressurization. The percentages of cells with Swi4-GFP in the nucleus in more than 300 cells are shown. (B) Mutant strains were cultured in SC medium at 0.1 or 25 MPa. Data are presented as mean ± standard deviation of three independent experiments.

### High pressure facilitates water influx into the cells

While conducting this study, we noticed that the wild-type cells were moderately swollen after incubation at 25 MPa. To validate this further, we examined whether the distribution of eisosomes was affected by pressure. Eisosomes, peripheral membrane protein complexes located on the cytoplasmic side of membrane compartments occupied by Can1 (MCC), are membrane invaginations (Douglas and Konopka, 2014; Lanze *et al.*, 2020; Sakata *et al.*, 2022). If the cell swells and the plasma membrane undergoes stretching under high pressure, the membrane invaginations should disappear. To verify this, we observed cells co-expressing the low-affinity tryptophan permease Tat1-GFP as a plasma membrane marker (Ishii *et al.*, 2022; Suzuki *et al.*, 2013) and an eisosome resident protein, Nce102-mCherry, after cultured at 25 or 50 MPa. Cells were observed at 0.1 MPa within 15 min of depressurization. At 0.1 MPa, Nce102-mCherry was localized to the plasma membrane in a patchy pattern characteristic of eisosomes (Fig. 5A). Notably, after high-pressure culture at 25 or 50 MPa, Nce102-mCherry was evenly distributed within the plasma membrane (Fig. 5A). This was partially suppressed by the presence of 1 M sorbitol (Fig. 5A). When the co-localization of Tat1-GFP and Nce102-mCherry was examined at the cell surface, Pearson’s correlation coefficient (PCC) increased with incubation time at 25 MPa (Fig. 5B), confirming the even distribution of Nce102 under high pressure. Similar increases in PCC with Tat1 were found for Pil1 and Sur7, another two eisosome-associated proteins, but not as pronounced as for Nce102 (Fig. 5B). These results suggest that the influx of water into the cells after high-pressure incubation results in tension in the plasma membrane. In the *wsc1*Δ and *slt2*Δ mutants, Nce102-mCherry was not localized in a patchy pattern but was more evenly distributed within the plasma membrane even under 0.1 MPa, suggesting excess water influx (data not shown).

**Fig. 5.**
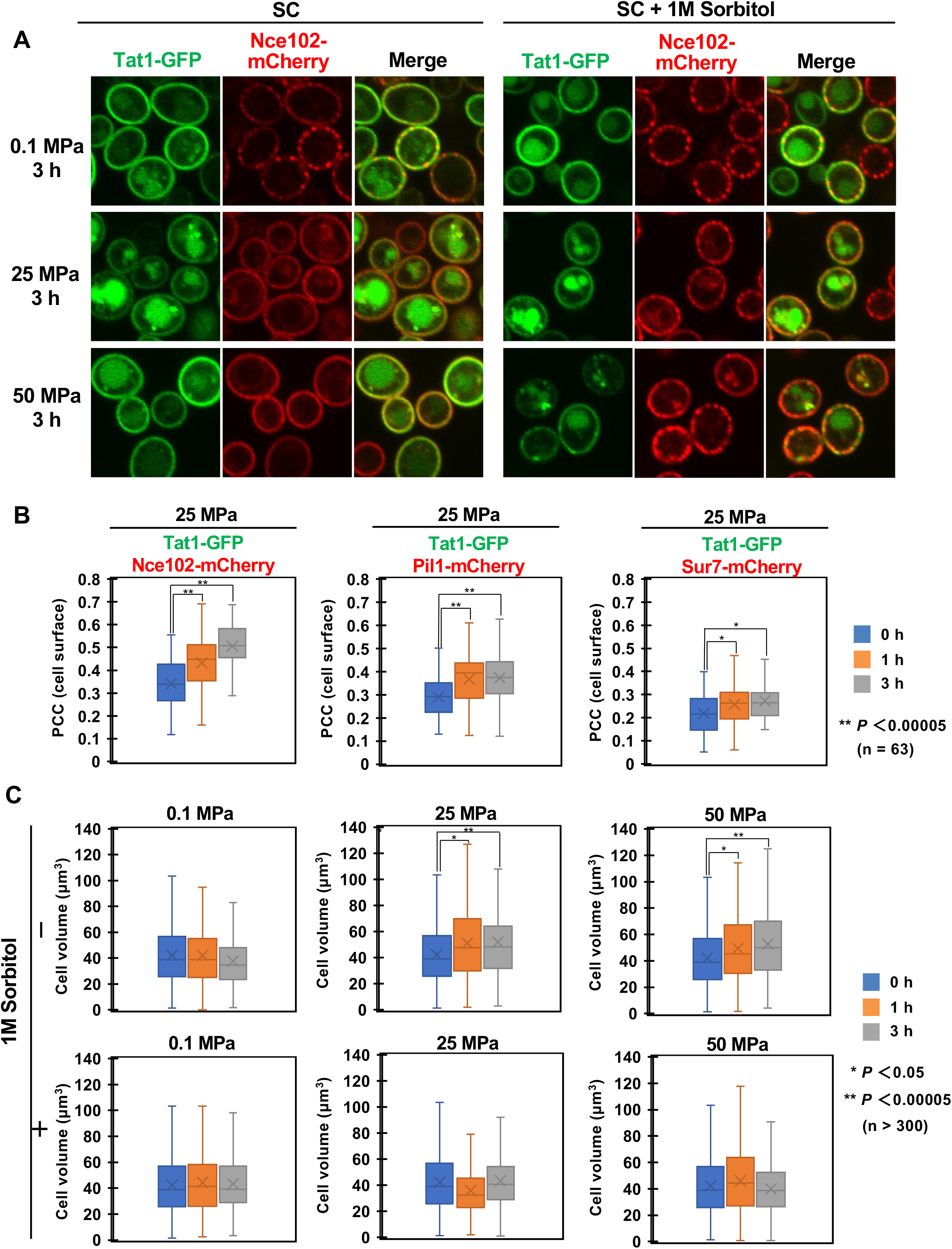
High pressure causes cell swelling and loss of the eisosome structure. (A) Wild-type cells co-expressing Tat1-GFP (plasma membrane marker) and Nce102-mCherry (eisosome marker) were cultured in SC medium at 0.1, 25, or 50 MPa for 3 h. After depressurization, cells were observed at 0.1 MPa under a confocal laser microscope. Cross-section images are shown. (B) Cells co-expressing Tat1-GFP and Nce102-mCherry, Pil1-mCherry, or Sur7-mCherry were cultured in SC medium at 0.1 or 25 MPa for 1 or 3 h. Pearson’s correlation coefficient (PCC) was obtained for cell surface images from 63 cells in three independent experiments. (C) Changes in cell volume when cultured under high pressure. Cells were cultured at 0.1, 25, or 50 MPa for 1 or 3 h in SC medium with or without 1 M sorbitol. The cell volume was calculated from the cell diameter of more than 300 cells after depressurization in three independent experiments.

Next, the cell volume was calculated by measuring the cell diameter using Tat1-GFP as the plasma membrane marker. We found that the cell volume increased 1.2-fold after incubation at 25 MPa or 50 MPa for 1 or 3 h, suggesting that water influx occurred under high pressure (Fig. 5A, B). To verify pressure-induced cell swelling, we examined the effects of 1 M sorbitol addition on cell volume under high pressure. As expected, 1 M sorbitol suppressed the high pressure-induced increase in cell volume (Fig. 5C). If high pressure merely pushed water into cells, the cell size would be restored following depressurization. However, the above results show that the cells remain swollen 15 min post depressurization until microscopic observation at atmospheric pressure. Therefore, even 1 h of incubation at 25 or 50 MPa would have caused irreversible changes to the cell wall. However, it remains unclear whether a high-pressure load directly weakens the cell wall structure or the cell wall expanded irreversibly due to increased water influx into the cell due to high-pressure application. In any case, cells were at higher risk of rupture under high pressure; therefore, to cope with this, cells tend to prevent water influx by expelling glycerol, the major osmolyte in yeast, to reduce internal osmotic pressure. Therefore, we focused on the regulation of intracellular glycerol levels under high hydrostatic pressure.

### Aquaglyceroporin Fps1-mediated glycerol efflux is required for high-pressure growth

Under hyperosmotic conditions, yeast cells synthesize or retain high concentrations of glycerol to maintain their osmotic balance (Blomberg, 2022; de Nadal and Posas, 2022; Hohmann, 2002). Intracellular glycerol concentration is partially regulated by Fps1, an aquaglyceroporin on the plasma membrane. Fps1 closes when extracellular osmolarity is high and opens when it is low, to expel glycerol and prevent cell rupture (Ahmadpour *et al.*, 2016; Beese *et al.*, 2009; Lee *et al.*, 2013; Luyten *et al.*, 1995; Muir *et al.*, 2015). The growth of the *fps1*Δ mutant was sensitive at 25 MPa (Fig. 6A). The addition of 1 M sorbitol to the medium restored the growth of the *fps1*Δ mutant to the wild-type level, at 25 MPa (Fig. 6A). Disruption of the glycerol synthase genes *GPP1* or *GPD1* increased, albeit partially, the high-pressure growth ability of the *fps1*Δ mutant (Fig. 5C). Furthermore, single deletion of *GPP1* enhanced high-pressure growth in the wild-type strain, although deletion of other glycerol biosynthetic genes had no promotive effect (Fig. 6B).

**Fig. 6.**
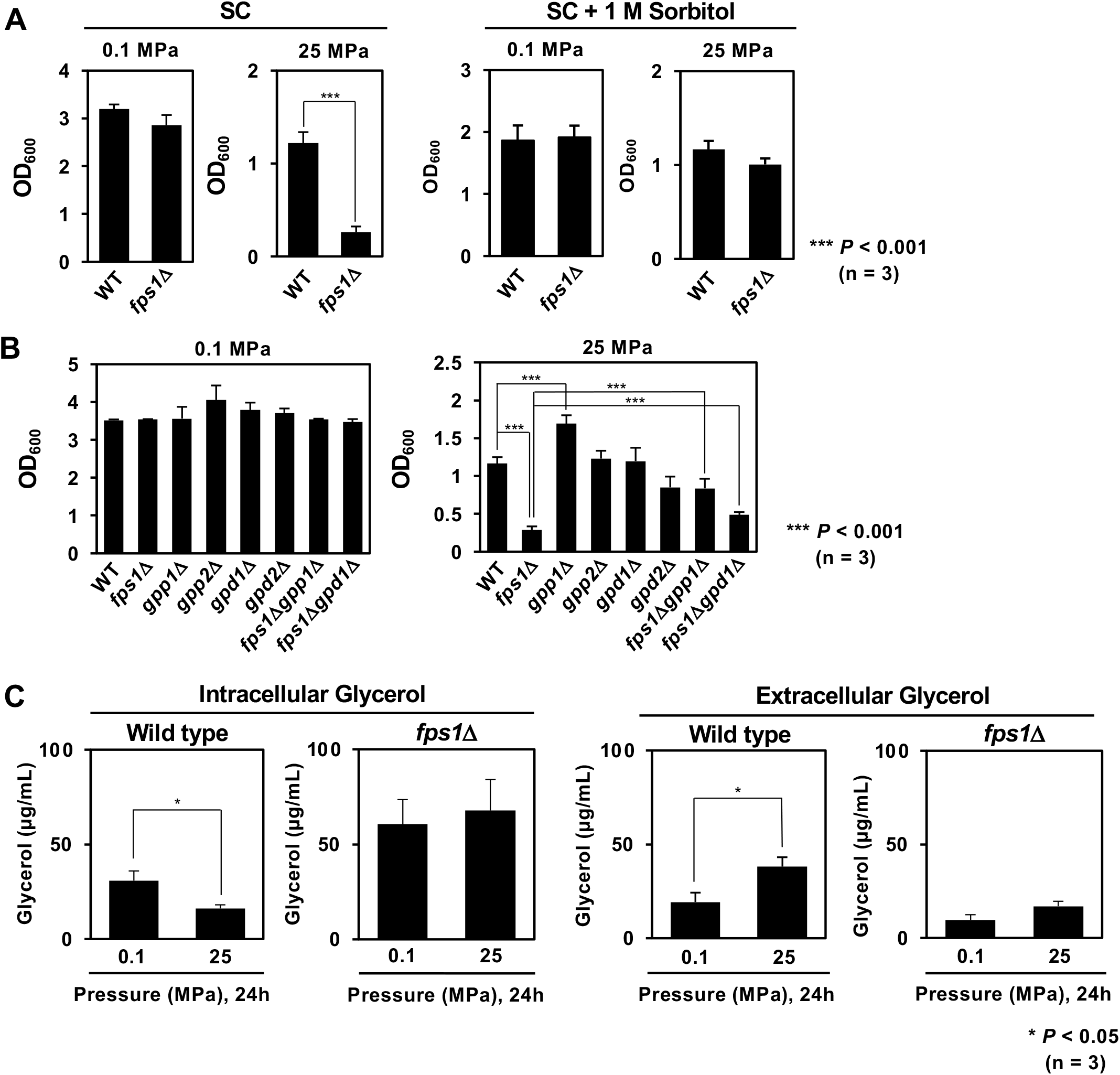
Glycerol efflux via Aquaglyceroporin Fps1 is required for high-pressure growth. (A) Wild-type and *fps1*Δ cells were cultured in SC medium with or without 1 M sorbitol at 0.1 or 25 MPa. (B) Mutants defective in glycerol synthesis were cultured in SC medium at 0.1 or 25 MPa. (C) Measurement of intracellular and extracellular glycerol levels after incubating cells at 0.1 or 25 MPa for 24 h with an initial OD_600_ = 0.1. Glycerol levels were quantified using a Glycerol assay kit. Data are presented as mean ± standard deviation of three independent experiments.

Next, we analyzed the intracellular and extracellular glycerol concentrations. Wild-type and *fps1*Δ cells in SC medium were incubated at 0.1 MPa and 25 MPa for 24 h, starting with an initial OD_600_ of 0.1. Upon analyzing these cells, we found that the intracellular glycerol concentration in wild-type cells decreased markedly after high-pressure culture, whereas more glycerol accumulated extracellularly (Fig. 6C). By contrast, *fps1*Δ cells had a 3-fold higher intracellular glycerol concentration than wild-type cells, which remained unchanged even after high-pressure culture (Fig. 6C). Thus, we concluded that wild-type cells expelled glycerol via Fps1 to cope with the high-pressure-induced excess water influx.

Furthermore, the ultrastructure of cells via transmission electron microscopy (TEM) showed that wild-type cells underwent no significant changes in cell morphology or cell wall structure after incubation at 25 MPa (Fig. 7). The ultrastructure of *fps1*Δ cells also appeared normal at 0.1 MPa. Interestingly, 20–40% of *fps1*Δ cells had sharply ruptured cell walls, without any abnormalities in their intracellular structures at 25 MPa (Fig. 7, arrows). The *fps1*Δ cell wall is weakened by increased turgor pressure due to glycerol accumulation; high pressure might also have caused further damage to the cell wall, leading to cell rupture.

**Fig. 7.**
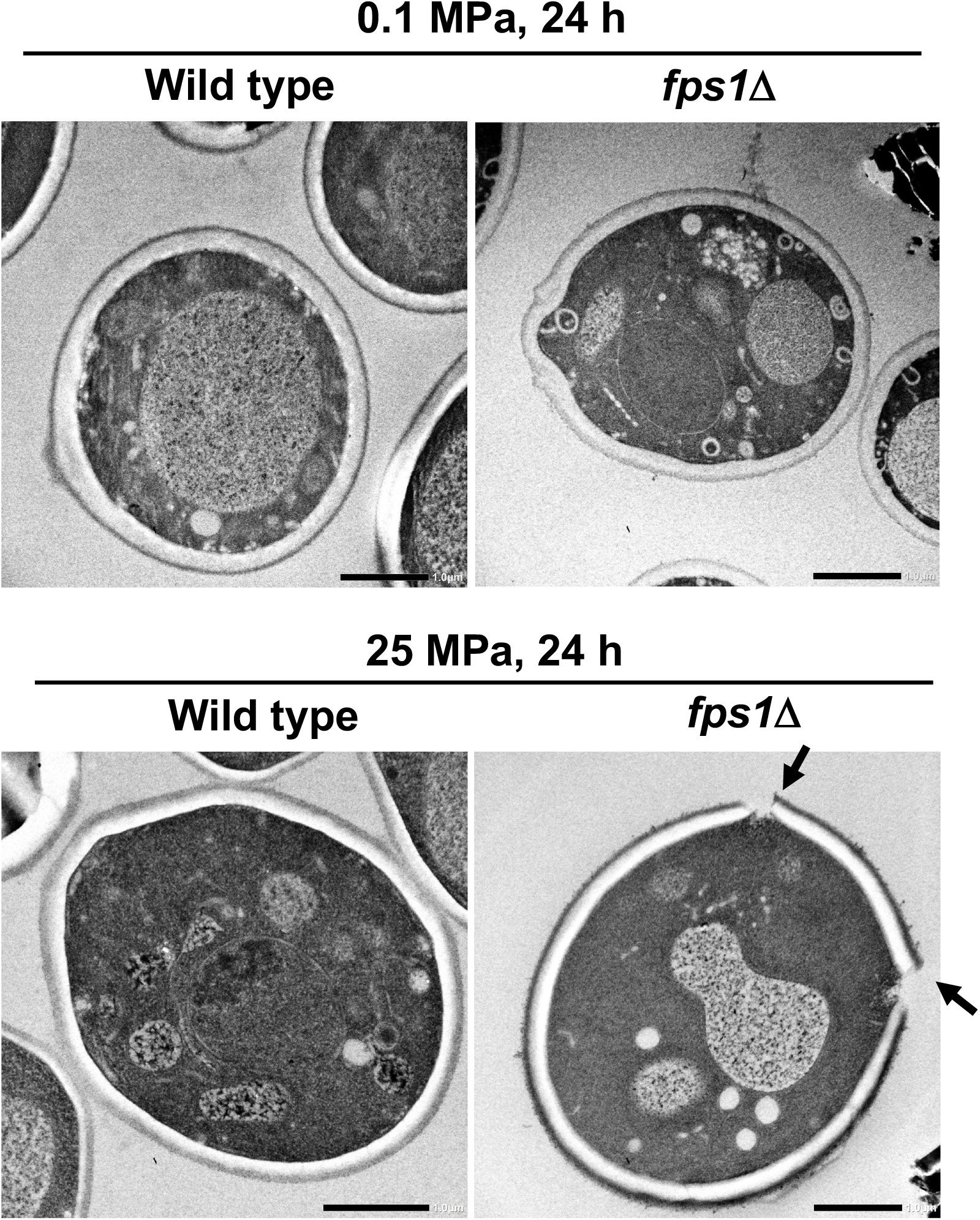
Ultrastructure of the cells observed using transmission electron microscopy. Wild-type and *fps1*Δ cells were cultured at 0.1 or 25 MPa for 24 h, and after depressurization, the samples were subjected to rapid freezing using the sandwich method. In *fps1*Δ cells, the points of cell wall breakage are indicated by arrows. Scale bars, 1.0 µm

### Fps1 opening is likely regulated by Slt2 during high-pressure cultures

The upstream regulators of Fps1 were analyzed to determine the mechanism by which cells regulate Fps1 in response to high pressure. Previous studies have indicated that Hog1 is a negative regulator of Fps1 opening, whereas Ypk1 and Slt2 are positive regulators (Fig. 1)(Ahmadpour *et al.*, 2016; Lee *et al.*, 2013; Muir *et al.*, 2015). To determine which kinases are involved in the regulation of the Fps1 in response to high pressure, we examined the high-pressure growth ability of the *fps1*Δ mutant expressing Fps1-T231A (Hog1 target site) or S537A (facilitates binding with Slt2) at 25 MPa. We found that the Fps1-S537A strain showed high sensitivity, whereas the Fps1-T231A strain, as well as the wild-type strain, grew at 25 MPa (Fig. 8A). We examined the phosphorylation of Fps1 using western blotting and observed a high-molecular-weight band of Fps1-3HA, the intensity of which increased slightly (2.2 ± 0.3-fold, n = 3) with increasing pressure at 25 MPa (Fig. 8B). As expected, the high molecular weight band of Fps1-3HA disappeared in cells expressing Fps1-S537A (Fig. 8B). Phosphorylation of the S537 residue of Fps1 is essential for its interaction with Slt2; however, there is no evidence of its direct phosphorylation by Slt2 (Ahmadpour *et al.*, 2016).

**Fig. 8.**
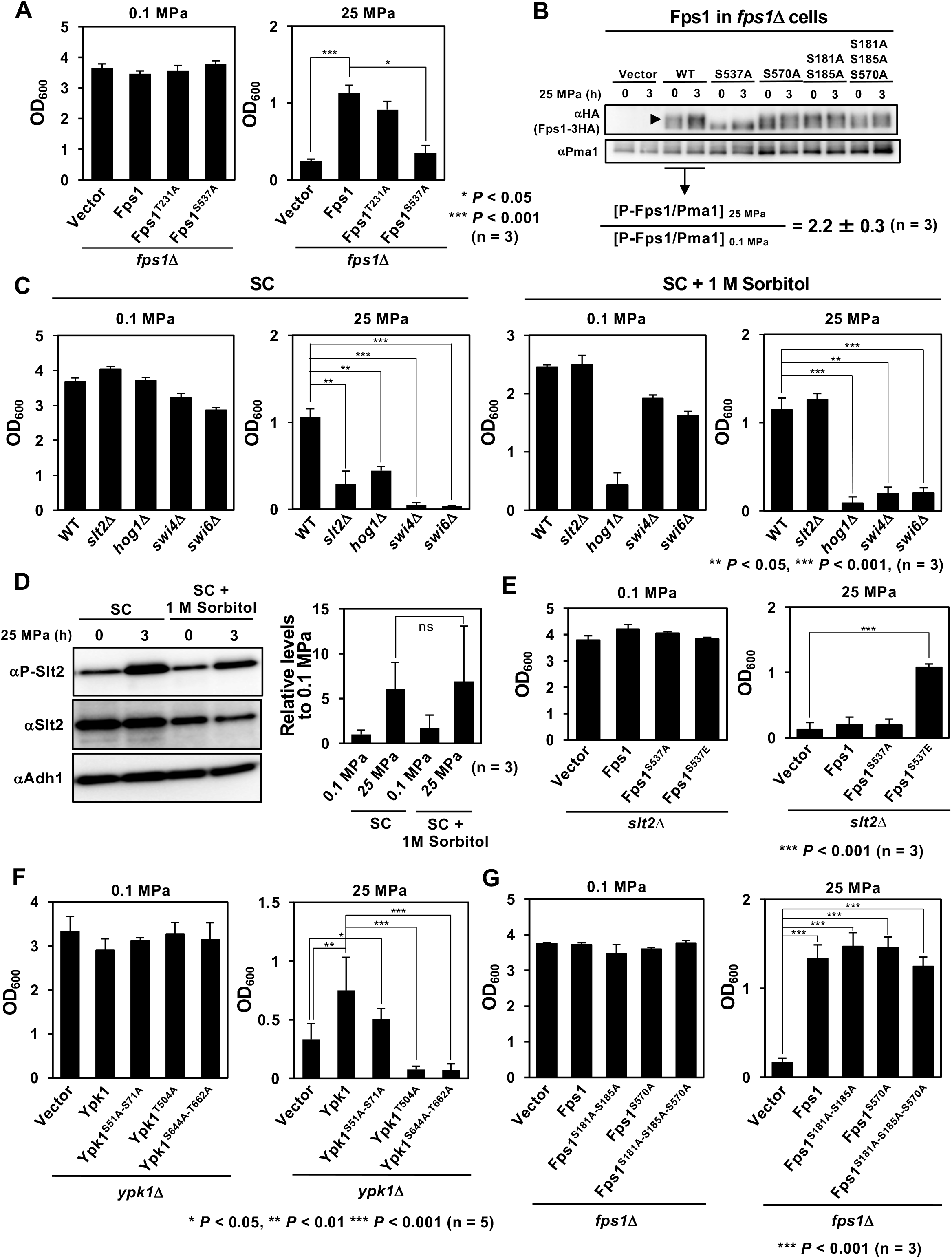
Slt2-dependent phosphorylation of Fps1 is required for growth under high pressure. (A) The Fps1 phosphorylation-site mutants were cultured in SC medium at 0.1 or 25 MPa. (B) Electrophoretic mobility of Fps1-3HA and its mutant forms was analyzed using western blotting. Cells were cultured at 0.1 or 25 MPa for 3 h, and whole cell extracts were prepared. The arrowhead indicates a higher-molecular weight band of Phospho-Fps1 (P-Fps1). The relative intensities, P-Fps1/Pma1 at 0.1 and 25 MPa, were quantified in an ImageQuant LAS4000 mini. (C) Wild-type, *slt2*Δ, *hog1*Δ, *swi4*Δ, and *swi6*Δ cells were cultured in SC medium with or without 1 M sorbitol at 0.1 or 25 MPa. (D) Wild-type cells were cultured in SC medium with or without 1 M sorbitol at 0.1 or 25 MPa for 3 h. Western blotting was performed as in Fig. 3D. (E) *slt2*Δ cells expressing Fps1, Fps1^S537A^ or Fps1^S537E^ were cultured in SC medium at 0.1 or 25 MPa. (F) The Ypk1 phosphorylation-site mutants were cultured in SC medium at 0.1 or 25 MPa. (G) The Fps1 phosphorylation-site mutants were cultured in SC medium at 0.1 or 25 MPa. Data are presented as mean ± standard deviation of three independent experiments.

The *slt2*Δ mutant was particularly sensitive to high pressure (Fig. 3C, 8C), which was completely rescued by the addition of 1M sorbitol (Fig. 8C). Therefore, the high-pressure sensitivity of the *slt2*Δ mutant was attributed to the excess water influx, at least in part, owing to its inability to open Fps1 and expel glycerol. The *hog1*Δ mutant also showed high-pressure sensitivity although the Fps1-T231A mutant grew at 25 MPa (Fig. 8A). The result suggests that Hog1 plays a role other than the Fps1 regulation under high pressure (Fig. 8C). The addition of 1M sorbitol did not restore high-pressure growth ability of the *hog1*Δ, *swi4*Δ, and *swi6*Δ mutants (Fig. 8C). Interestingly, Slt2 was also hyperphosphorylated when cultured with 1 M sorbitol at 25 MPa where there is no excessive influx of water into cells (Fig. 8D). The result suggests that the activation of the CWI pathway by high pressure may also be caused by a direct effect of high pressure on Wsc1, in addition to its activation via the cell swelling associated with the excessive influx of water under high pressure.

To verify whether the high-pressure sensitivity of the *slt2*Δ mutant was due to reduced Fps1 activity, we expressed Fps1^S537E^, a simulated phosphorylated form of Fps1 at the S537 residue, and found that it efficiently rescued the high-pressure sensitivity of the *slt2*Δ mutant (Fig. 8E). Therefore, Slt2 activates Fps1, thereby preventing water influx and allowing for cell growth under high pressure. Given that 1 M sorbitol fully restored the high-pressure growth ability of the *slt2*Δ mutant (Fig. 8C), it is possible that Slt2 may also prevent water influx under high pressure by another mechanism, in addition to activating Fps1.

### Ypk1 is required for growth under high pressure, independent of the regulation of Fps1

Fps1 is also phosphorylated by Ypk1 for its activation, depending on TORC2 signaling (Muir *et al.*, 2015). We found that the disruption of *YPK1* caused a severe defect in cell growth at 25 MPa, demonstrating the importance of Ypk1 (Fig. 8F). Ypk1 is phosphorylated by multiple protein kinases such as Fpk1/Fpk2 (Rispal *et al.*, 2015), Pkh1/Pkh2 (Roelants *et al.*, 2002; Roelants *et al.*, 2004; Roelants *et al.*, 2010), and TORC2 (Leskoske *et al.*, 2017; Muir *et al.*, 2015; Roelants *et al.*, 2004) (Fig. 1). To infer which kinases are required for high-pressure growth via phosphorylation of Ypk1, we examined the effect of alanine substitutions for S51/S57 residues (Fpk1/Fpk2 target sites for Ypk1 inactivation), T504 residue (Pkh1/Pkh2 target site for Ypk1 activation), and S644/T662 residues (TORC2 target sites for Ypk1 activation) on cell growth under high pressure (Fig. 1). While cells expressing Ypk1-S51A-S71A grew as efficiently as wild-type cells at 25 MPa, those expressing Ypk1-T504A or S644A-T662A showed severe growth defects under the pressure (Fig. 8F). Thus, it is possible that Pkh1/Pkh2 and/or TORC2 phosphorylate Ypk1 in response to high pressure. As shown in Fig. 4, cells swelled after high-pressure incubation, and the eisosome structure disappeared. This raises the possibility that Slm1, localized in the eisosome, is released by high pressure and eventually binds to and activates TORC2, ultimately phosphorylating Ypk1 (Fig. 1)(Berchtold *et al.*, 2012; Riggi *et al.*, 2018). To test this hypothesis in our preliminary experiments, Lsp1-mCherry/Slm1-GFP-co-expressing and Avo3-3×mCherry/Slm1-GFP-co-expressing strains were constructed, and any alterations in the colocalization of each strain under high pressure were observed using confocal laser microscopy. Avo3 and Lsp1 are components of TORC2 and the eisosome, respectively (Babst, 2019; Eltschinger and Loewith, 2016). We found that the colocalization of Lsp1-mCherry and Slm1-GFP decreased after cell incubation at 25 MPa, whereas that of Avo3-3×mCherry and Slm1-GFP appeared to increase, suggesting TORC2 activation (Fig. S1A). Furthermore, phospho-Ypk1 was detected using a rabbit monoclonal antibody for phospho-PKC (pan) (zeta Thr410) (190D10), which reacts with the phosphorylated T504 residue, and after incubation at 25 MPa for 3 h, its phosphorylation level increased 7-fold. (Fig. S1B).

To test whether phosphorylation of Fps1 by Ypk1 is required for high-pressure growth, the effect of phosphorylation site mutations was examined. Residue S570 of Fps1 is majorly phosphorylated by Ypk1, while the phosphorylation of residues S181 and S185 has also been reported (Fig. 1) (Muir *et al.*, 2015). Both the Fps1-S570A-substituted and S181A-S185A-S570A triple-substituted strains grew as well as the wild-type strain (Fig. 8G). Consistent with this result, the electrophoretic mobility of the alanine substitution mutants Fps1-S570A-3HA, Fps1-A181A-S185A-3HA, and Fps1-A181A-S185A-S570A-3HA remained unaffected (Fig. 8B). Thus, Ypk1 is unlikely to phosphorylate Fps1 in response to high pressure.

## Discussion

Based on the coincidental observation that cells moderately swell under high pressure, the importance of glycerol efflux by Fps1 and its regulation via the CWI pathway has been the focus of this study. We propose the presence of high-pressure signaling systems mediated by the stress sensor Wsc1 (Fig. 9). In our model, high pressure activates the CWI pathway in two distinct modes. First, high pressure promotes water influx into yeast cells, thereby increasing turgor pressure, which in turn expands the cell walls and activates Wsc1 (Fig. 2–5). Second, high pressure directly activates Wsc1, as suggested by our finding that Slt2 was hyperphosphorylated at 25 MPa even in the absence of cell swelling in 1 M sorbitol medium (Fig. 8D). From the first perspective, the water influx caused by increased hydrostatic pressure is similar to that of hypoosmotic stress. In growing yeast cells, turgor pressure is estimated to be 0.5–1.5 MPa due to the influx of water driven by the osmotic gradient (Atilgan *et al.*, 2015; Minc *et al.*, 2014; Mishra *et al.*, 2022; Schaber *et al.*, 2010). The turgor pushes the cell membrane, further expanding the cell wall. Using an osmometer, we found the osmotic pressure of the SC medium to be 255 ± 0 mOsm/kg (n = 3), which corresponds to 0.588 MPa (nearly 0.6 MPa). Assuming that the turgor pressure of the cells was 1.0 MPa, the intracellular osmotic pressure was 1.6 MPa. Applying hydrostatic pressure to the cell culture caused a water influx into the cells until the water potentials inside and outside the cell became equivalent. Conversely, from the second perspective, it is possible that high pressure of 25 MPa directly modulates Wsc1 structure or promotes physical interaction between Wsc1 and Rom2; however, evidence of high pressure-induced structural changes in Wsc1 or enhanced Wsc1–Rom2 interaction under high pressure remains lacking.

**Fig. 9.**
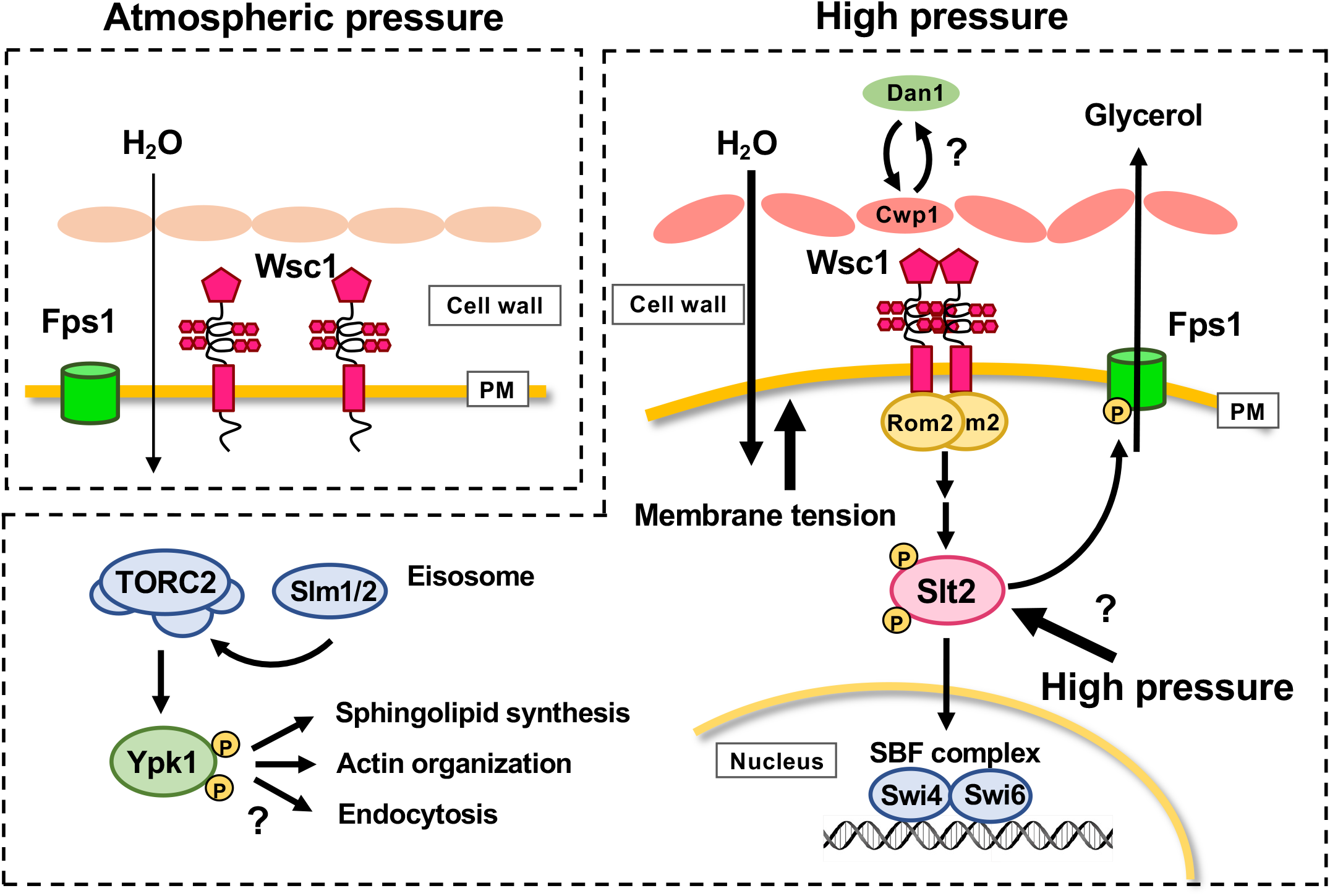
Model depicting Wsc1-dependent high-pressure signaling pathway in yeast. When pressure is applied to the cell, an excessive influx of water into the cell occurs. Perturbation of the cell wall by high pressure causes further water influx, and the cell becomes at a risk of rupture. To prevent this, Wsc1 detects cell wall stretching and activates the CWI pathway. Once the CWI pathway is activated, Slt2 kinase phosphorylates the aquaglyceroporin Fps1. Phosphorylated Fps1 promotes its opening, which allows cells to expel glycerol, adjusting the osmotic pressure balance across the plasma membrane. TORC2-Ypk1 pathway is independent of the regulation of Fps1 under high pressure.

High pressure may also cause some perturbation to the cell wall, as suggested by our observations that cells remain swollen at least 15 min post depressurization (Fig. 5). However, there is currently no direct evidence that high pressures of 25 MPa damage cell walls. Since high hydrostatic pressure has little effect on covalent binding (Chen *et al.*, 2017), it is unlikely that the β1,3-glucan or β1,6-glucan networks are cleaved by high pressure. Rather, we suspect that pressure may have a greater effect on the levels of cell wall mannoproteins. Our previous DNA microarray results showed that transcription of the gene encoding for the major cell wall mannoprotein Cwp1 was markedly reduced under 25 MPa; instead, the *DAN/TIR* family mannoprotein genes, which are typically induced under hypoxic conditions (Abramova *et al.*, 2001), were significantly enhanced (Abe, 2007). Although their protein levels have not been examined, the composition of mannoproteins may change with high pressure, and eventually, cell wall elasticity may change. Once the CWI pathway is activated, Slt2 kinase promotes glycerol efflux through aquaglyceroporin Fps1, thereby allowing for cells to expel glycerol and adjust the osmotic pressure balance across the plasma membrane (Fig. 6). Under high pressure, the high molecular weight band of Fps1 increased, suggesting enhanced Fps1 phosphorylation. However, a similar high molecular weight band of Fps1 was also observed in the *slt2*Δ mutant, although it disappeared in Fps1^S537A^. This suggests other kinases besides Slt2 that phosphorylate the Fps1 S537 residue. Since phosphorylation of Fps1 S537 residues is critical for binding to Slt2, there may be other sites of authentic phosphorylation by Slt2. The role of Slt2 in growth under high pressure is not only limited to the activation of Fps1, but also the phosphorylation of Swi4/Swi6 and the induction of transcription of many downstream genes. This is justified by the fact that loss of *SWI4* or *SWI6* causes severe high-pressure sensitivity (Fig. 4).

As per the TEM images of *fps1*Δ cells, the cell wall ruptured sharply after a high-pressure incubation (Fig. 7). This observation may be due to the localized weakening of the cell wall by increased turgor pressure due to glycerol accumulation in the *fps1*Δ cells. High pressure may have caused further damage to the cell wall in the *fps1*Δ mutant, leading to cell rupture. However, we cannot rule out the possibility that the weakened *fps1*Δ cell walls were ruptured during TEM sample preparation by the sandwich freezing method (Yamaguchi *et al.*, 2021a; Yamaguchi *et al.*, 2021b).

Although this study showed the importance of Ypk1 kinase in high-pressure growth (Fig. 8F), it did not provide evidence that Ypk1 regulates the opening of Fps1 by phosphorylation (Fig. 8G). Rather, alanine substitutions at Ypk1 residues, phosphorylated by TORC2 and Pkh1/Pkh2 kinases, prevented high-pressure growth (Fig. 8F). TORC2-Ypk1 regulates polarization of the actin cytoskeleton and sphingolipid biosynthesis (Kamada *et al.*, 2005; Rispal *et al.*, 2015; Roelants *et al.*, 2011; Tabuchi *et al.*, 2006). Therefore, the regulation of sphingolipid synthesis or actin organization may be important for high-pressure adaptation. Indeed, high pressures stiffen membrane lipid bilayers (Matsuki *et al.*, 2013; Winter, 2002), which may adversely affect membrane protein functions, membrane fusion, or membrane trafficking. Furthermore, hypoosmotic stress causes the depolymerization of the actin cytoskeleton (Gualtieri *et al.*, 2004). To cope with these pleiotropic effects of high pressure, yeast cells may activate Fps1 via the CWI pathway to expel glycerol and avoid cell rupture, promote sphingolipid synthesis via the TORC2-Ypk1 pathway to maintain membrane homeostasis, and regulate actin cytoskeleton organization.

Therefore, the regulation of sphingolipid synthesis and actin cytoskeleton via the TORC2-Ypk1 pathway is an interesting research target for understanding the physiological responses to high hydrostatic pressure in yeast.

Our previous studies have shown that defects in *TRP1*, *TAT2*, TORC1-EGOC, or poorly characterized genes such as *EHG1/MAY24* cause severe high-pressure sensitivity (Abe and Horikoshi, 2000; Abe and Minegishi, 2008; Kurosaka *et al.*, 2019; Uemura *et al.*, 2020). On the contrary, the defects in the CWI pathway and Fps1 shown in this study caused relatively mild sensitivity to high-pressure. This is probably because the former defects cause impaired nutrient uptake or aberrant glutamine accumulation under 25 MPa, resulting in rapid growth arrest after pressurization, whereas the latter defects are due to an inability to prevent water influx under high pressure, which causes the negative effects on growth to progress slowly. Yeasts possess two types of aquaporins, Aqy1 and Aqy2. However, both alleles are nonfunctional in experimental strains derived from S288C, such as strain BY4742 (Sabir *et al.*, 2017). Therefore, the water influx into the cell when exposed to hypoosmotic or high hydrostatic pressure stress is thought to be mainly due to passive diffusion across the plasma membrane lipid bilayers. Passive diffusion of water across the lipid bilayer is much slower than water flux through aquaporins. This could explain why the effects of high-pressure stress are relatively mild. The Σ1278b strain and the naturally occurring *S. cerevisiae* strains have functional *AQY* alleles (Laize *et al.*, 1999). Therefore, the effects of hypoosmotic and high-pressure stress in these strains could be more severe than in the experimental strains. The osmotic pressure of the SC medium was 255 mOsm/kg, which corresponds to 127.5 mM NaCl. When it rains, the ambient osmotic pressure drops to near zero, exposing naturally occurring yeasts to greater hypoosmotic stress. Exposure to high hydrostatic pressure under such low osmotic pressure would pose a threat to their survival.

Meanwhile, the ambient osmotic pressure is isotonic for human and other animal cells. Therefore, excessive water influx due to high water pressure loading, as observed in yeast in this study, would not occur. However, when human endothelial cells are subjected to a pressure of 50 mmHg (~6.7 kPa), which corresponds to a transient increase in blood pressure during exercise, water is extruded from the cell via aquaporin 1, causing cell contraction and plasma membrane deformation (Yoshino *et al.*, 2020). Consequently, protein kinase C is activated via the G protein-coupled receptor and phospholipase C, which in turn activates the Ras/ERK pathway, ultimately leading to the formation of complex tubular vascular structures (Yoshino *et al.*, 2020).

Since aquaporins are channels, if the water potential is equivalent inside and outside the cells, no water efflux should occur even when the pressure is 50 mmHg. However, because the pressure is very weak, the pressure applied at different sites in a single cell may be different, and water may be extruded through aquaporins at lower-pressure sites.

The effects of high hydrostatic pressure vary not only with the magnitude of the applied pressure but also with the duration of the pressurization. Therefore, the effects on the complex metabolic processes of living cell systems cannot be generalized. The full picture of pressure effects will be revealed by applying the extensive knowledge of yeast cell biology and sophisticated molecular genetic tools to advanced transcriptomics, proteomics, and metabolomics studies, which could potentially be translated to mammalian cells and help improve cellular health under normal and stress conditions.

## Materials and methods

### Yeast strains and media

The parental wild-type strain BY4742 and the deletion mutants used in this study are listed in Table 1. Strains DL376 (*pkc1*Δ) and DL100 (the parental strain of DL376) were kindly provided by Dr. David Levin of Boston University Goldman School of Dental Medicine through Dr. Hidetoshi Iida of Tokyo Gakugei University (Levin and Bartlett-Heubusch, 1992). The two strains were transformed with YCplac22 (*TRP1, CEN*) to provide tryptophan prototrophy. Cells were grown in SC medium or YPD medium at 25 °C under 0.1 MPa or 25 MPa (Abe and Minegishi, 2008). The SC medium contained 0.67% yeast nitrogen base without amino acids (Becton, Dickinson and Company, MD, USA), 2% D-glucose, 20 μg/mL methionine, 90 μg/mL leucine, 30 μg/mL isoleucine, 40 μg/mL tryptophan, 20 μg/mL histidine, 30 μg/mL lysine, 20 μg/mL methionine, 50 μg/mL phenylalanine, 30 μg/mL tyrosine, 20 μg/mL arginine, 100 μg/mL aspartic acid, 100 μg/mL glutamic acid, 400 μg/mL serine, and 200 μg/mL threonine. YPD medium contains 1% Bacto yeast extract (Becton, Dickinson and Company), 2% Bacto peptone (Becton, Dickinson and Company), and 2% D-glucose.

**Table 1.**
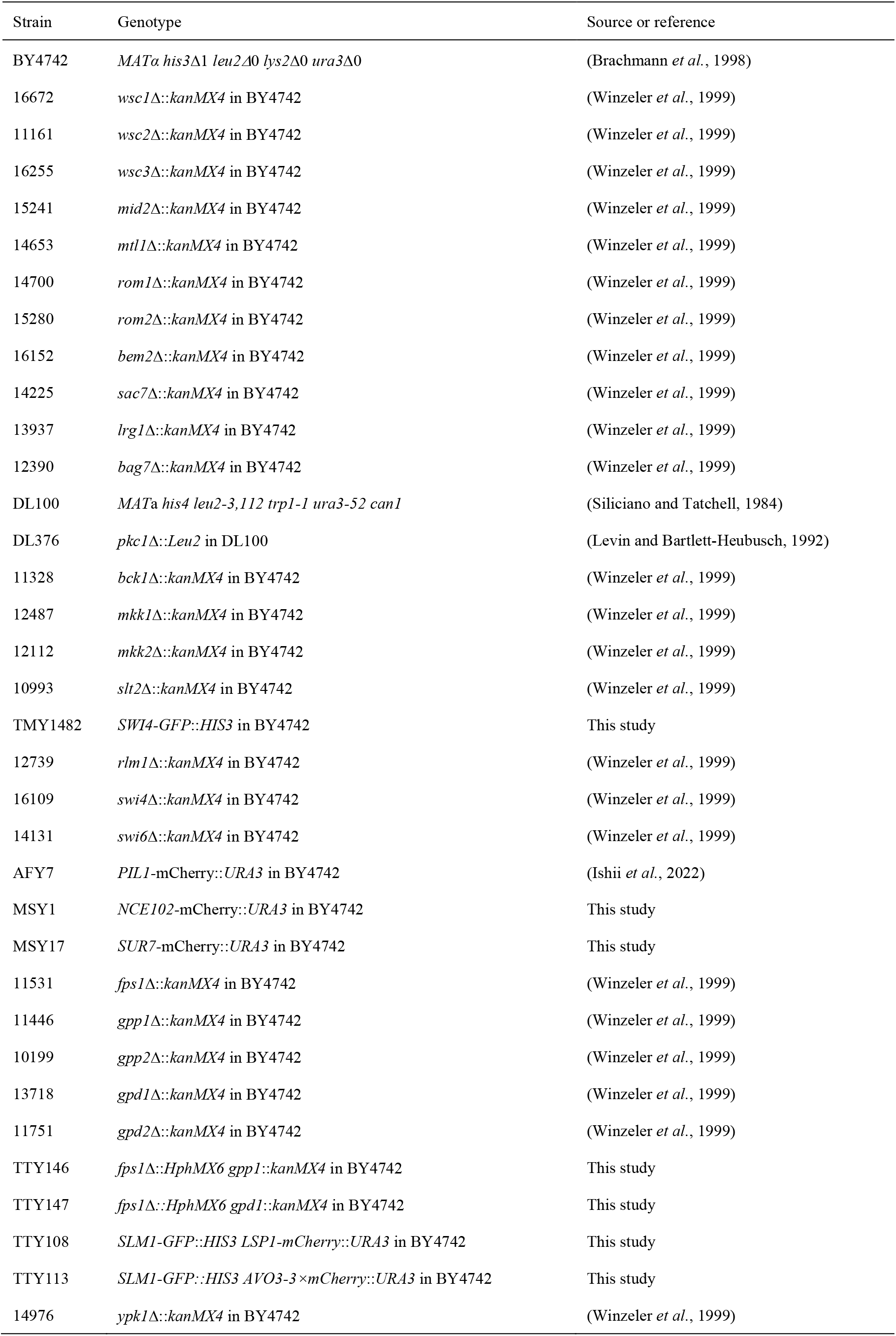
Strains used in this study

### High-pressure cell culture

For most experiments examining growth efficiency under high pressure, growing cells in shaking culture were diluted in SC or YPD medium to an OD_600_ of 0.01. The diluted cells were transferred to sterile 2.2-mL tubes and sealed with Parafilm^TM^ (Bemis Corp., Neenah, WI, USA). The tubes were placed in a 500-mL high-pressure chamber (Abe and Minegishi, 2008) and subjected to 25 MPa at 25 °C for 48 h. After depressurization, cell growth was evaluated by measuring OD_600_ using a PD-303 spectrophotometer (Apel, Kawaguchi, Japan). Here, OD_600_ = 1.0 corresponds to 1.65 × 10^7^ cells/mL in our device setup.

### Generation of plasmids and mutant strains

The plasmids used in the present study are listed in Table 2. Polymerase chain reaction (PCR)-based site-directed mutagenesis was performed using the appropriate primers. The primer sequences will be disclosed upon reasonable request. Homologous recombination, using pJT6 (GFP, *HIS3*) and pJT601 (mCherry, *URA3*), was used to tag genes in the yeast genome with GFP and mCherry, respectively (kind gifts from Dr. Jiro Toshima of Tokyo University of Science, (Kawada *et al.*, 2015)). Plasmids encoding HA-Rho1 (SP333; *LEU2, CEN*), HA-Rho1^F35L^ (SP335; *LEU2, CEN*), and HA-Rho1^Q68H^ (SP336; *LEU2, CEN*) were kindly provided by Dr. Satoshi Yoshida of Waseda University (Yoshida *et al.*, 2009). pBSK-*SFK1*-3×mCherry-*CaURA3* was kindly provided by Dr. Takuma Kishimoto of Hokkaido University Graduate School of Life Science (Kishimoto *et al.*, 2021). To generate Avo3-3×mCherry strain, a DNA fragment encoding a *C*-terminal domain of Avo3 was amplified by PCR and was inserted into pBSK-*SFK1*-3×mCherry-*CaURA3* of which *SFK1* gene had been removed. The resulting plasmid was linearized by digestion with *Eco*RV and introduced into the wild-type strain harboring *SLM1*-GFP in the genome.

**Table 2.** Plasmids used in this study

| Plasmid | Description | Source or reference |
| --- | --- | --- |
| pUA35 | 3HA *URA3 CEN4* | Laboratory stock |
| pTM35 | *GFP URA3 CEN4* | Laboratory stock |
| YEplac181 | *LEU2 2 μ* | (Gietz and Sugino, 1988) |
| YCplac22 | *TRP1 CEN4* | (Gietz and Sugino, 1988) |
| pRS313 | *HIS3 CEN6* | (Sikorski and Hieter, 1989) |
| pTAT1-GFP | *TAT1*-GFP driven by its own promoter in pRS313 | This study |
| pMB1 | *WSC1*-3HA driven by its own promoter in pUA35 | This study |
| pMB23 | *WSC1*-3HA-Y303A driven by its own promoter in pUA35 | This study |
| pMB74 | *WSC1*-3HA-N344A-P345A-F346A driven by its own promoter in pUA35 | This study |
| pSS21 | *WSC1*-GFP driven by its own promoter in pTM35 | This study |
| pMB80 | *WSC1*-3HA-N344A-P345A-F346A driven by its own promoter in pTM35 | This study |
| SP333 | HA-*RHO1* driven by its own promoter *LEU2 CEN* | (Yoshida *et al.*, 2009) |
| SP335 | HA-*RHO1*-F35L driven by its own promoter *LEU2 CEN* | (Yoshida *et al.*, 2009) |
| SP336 | HA-*RHO1*-Q68H driven by its own promoter *LEU2 CEN* | (Yoshida *et al.*, 2009) |
| pME29 | *PKC1* driven by its own promoter in YEplac181 | This study |
| pME30 | *PKC1*-R398P driven by its own promoter in YEplac181 | This study |
| pTNK5 | *FPS1* driven by its own promoter in pUA35 | This study |
| pTNK8 | *FPS1*-T231A driven by its own promoter in pUA35 | This study |
| pTNK9 | *FPS1*-S537A driven by its own promoter in pUA35 | This study |
| pTNK10 | *FPS1*-S570A driven by its own promoter in pUA35 | This study |
| pTNK22 | *FPS1*-S181A-S185A driven by its own promoter in pUA35 | This study |
| pTNK23 | *FPS1*-S181A-S185A-S570A driven by its own promoter in pUA35 | This study |
| pSS10 | *FPS1*-3HA driven by its own promoter in pUA35 | This study |
| pSS26 | *FPS1*-3HA-S537A driven by its own promoter in pUA35 | This study |
| pSS12 | *FPS1*-3HA-S570A driven by its own promoter in pUA35 | This study |
| pSS16 | *FPS1*-3HA-S181A-S185A driven by its own promoter in pUA35 | This study |
| pSS14 | *FPS1*-3HA-S181A-S185A-S570A driven by its own promoter in pUA35 | This study |
| pSS6 | *YPK1* driven by its own promoter in pUA35 | This study |
| pTNK11 | *YPK1*-S51A-S71A driven by its own promoter in pUA35 | This study |
| pTNK7 | *YPK1*-T504A driven by its own promoter in pUA35 | This study |
| pTNK14 | *YPK1*-S644A-T662A driven by its own promoter in pUA35 | This study |
| pTM102 | pBSK (+) -*SFK1*-3×mCherry | (Kishimoto *et al.*, 2021) |
| pJT6 | GFP *HIS3* | (Kawada *et al.*, 2015) |
| pJT601 | mCherry *URA3* | (Kawada *et al.*, 2015) |

### Western blotting

Whole-cell extract preparation was performed as previously described (Ishii *et al.*, 2022). Briefly, 1.65 × 10^8^ cells were collected by centrifugation; serially washed with 10 mM NaN_3_, 10 mM NaF, and lysis buffer A (50 mM Tris-HCl, 5 mM EDTA, 10 mM NaN_3_, and Complete™ EDTA-free tablet; Roche Diagnostics, Mannheim, Germany); disrupted with glass beads at 4 °C. Unbroken cells and debris were removed by centrifugation at 900 × *g* for 5 min, and purified lysates were treated with 5% (w/v) Sodium dodecyl sulfate (SDS) and 5% (v/v) 2-mercaptoethanol at 37 °C for 10 min. Whole-cell extract preparation to detect phospho-Slt2 was performed as previously described (Wilk *et al.*, 2010) with some modifications. Yeast cells amounting to 5 × 10^7^ in SC medium were treated with 255 mM NaOH–1% 2-mercaptoethanol on ice for 15 min, then 6% trichloroacetic acid (TCA) was added and cells were kept on ice for another 15 min. The cell pellet was collected after centrifugation at 15,000 × *g* for 5 min, washed with 1 M Tris-HCl (pH 6.8), and was treated with a sample buffer (65 mM Tris [pH6.8], 2% SDS, 10% Glycerol, 100 mM DTT, 0.002% BPB) to denature proteins at 95 °C for 5 min. The supernatant after centrifugation at 15,000 × *g* for 5 min was used for western blotting. Whole-cell extract preparation to detect phospho-Ypk1was performed as previously described (Roelants *et al.*, 2011) with some modifications. Yeast cells amounting to 1.65 × 10^7^ were collected by centrifugation at 6,000 × *g* for 1 min. After removing the supernatant, the cell pellet was frozen in liquid nitrogen and was treated with 150 µL of 1.85 M NaOH–7.4% 2-mercaptoethanol for 15 min on ice. Then, 150 µL of 50% TCA was added to the cell suspension to precipitate proteins on ice for 10 min. After removing the supernatant, the proteins were washed twice with acetone and were denatured with the sample buffer at 95 °C for 5 min. Proteins were separated by SDS-PAGE and transferred to Immobilon PVDF membranes (EMD Millipore, Billerica, MA, USA). The membrane was incubated for 1 h with antibodies. The following antibodies were used: a monoclonal antibody for HA (Medical & Biological Laboratories Co. Ltd., Nagoya, Japan), a rabbit polyclonal antibody for Adh1 (Rockland Immunochemicals, Inc., Gilbersville, PA, USA), a rabbit polyclonal antibody for Pma1 (Usami *et al.*, 2014), a mouse monoclonal antibody for Slt2 (cat. no. sc-133189, Santa Cruz Biotechnology, TX, USA), a rabbit monoclonal antibody phospho-p44/42 MAPK (Erk1/2, Thr202/Tyr204; cat no. 4376, Cell Signaling Technology, MA, USA) to detect phospho-Slt2, a rabbit polyclonal antibody for Ypk1 (a kind gift from Dr. Mitsuaki Tabuchi of Kagawa University), and a rabbit monoclonal antibody for Phospho-PKC (pan) (zeta Thr410) (190D10) (cat. no. 2060, Signaling Technology) to detect phospho-Ypk1. The membranes were washed and incubated for 30 min with horseradish peroxidase-conjugated goat anti-rabbit IgG (GE Healthcare Bio-Sciences, NJ, USA). Labeling was performed using an enhanced chemiluminescence (ECL) select kit (GE Healthcare Bio-Sciences). The chemiluminescence signals were detected in an ImageQuant LAS4000 mini (GE Healthcare Life Sciences, NJ, USA)

### Image collection and analyses on confocal laser microscope

Cells expressing GFP- or mCherry-tagged proteins were imaged using a confocal laser microscope (FV-3000) equipped with an objective lens UPLAPO60XOHR (NA 1.5) (Olympus, Co. Ltd., Tokyo, Japan). To determine the cell volume, cells were assumed to be spheres or ellipsoids. Using Tat1-GFP as a cell membrane marker (Ishii *et al.*, 2022), cross-sectional images of > 100 cells cultured under each condition were acquired. CellSens software (Olympus, Co. Ltd) was used to mark Tat1-GFP as a circle or ellipse, and the radius (*r* µm) or long (*a* µm) and short radii (*b* µm) were calculated. For spheres, the volume (µm^3^) was approximated by V = 4/3π\**r*^3^, and for ellipsoids, *V* = 4/3π\**a*\**b*^2^. Changes in cell volume due to high-pressure incubation were measured by acquiring images at 0.1 MPa, immediately after depressurization. Colocalization of Lsp1-mCherry with Slm1-GFP or Avo3-3×mCherry with Slm1-GFP was analyzed in more than 50 cells from three independent experiments using cellSens software and represented by the Pearson colocalization coefficient. An IX70 epifluorescence inverted microscope (Olympus, Co. Ltd.) was used for imaging.

### Determination of glycerol concentration

Culture medium containing cells of 1.0 OD_600_ (1.65 × 10^7^ cells) was centrifuged at 6,000 × *g* for 1 min to separate the supernatant and cell pellet. The cell pellet was dissolved in 100 µL of sterilized deionized water and the mixture was heated at 95 °C for 10 min. After cooling to room temperature, the mixture was centrifuged at 21,500 × *g* for 10 min to collect the supernatant. Glycerol concentrations in the samples were determined using a glycerol colorimetric assay kit (item no. 10010755, Cayman Chemical Company, MI, USA) with an appropriate dilution.

### Measurement of osmotic pressure

The osmotic pressure of SC medium was measured using a Fiske 210 osmometer (Advanced Instruments, MA, USA).

### Transmission electron microscopy observation

Samples for TEM observation were prepared as described previously (Yamaguchi *et al.*, 2021a; Yamaguchi *et al.*, 2021b), with modifications. Cells cultured at 0.1 MPa or 25 MPa for 24 h in the SC medium were briefly centrifuged at 760 × *g* following depressurization, and less than 1 µL of the cell suspension was sandwiched between two copper disks (3 mm diameter) that had been prehydrophilized. The copper disks were then sandwiched between reverse-acting tweezers and dropped into a coolant for quick freezing. An isopentane: ethanol (4 : 1) mixture cooled with liquid nitrogen to a sherbet-like consistency (approximately −130 °C) was used as the coolant. The copper disks were immediately transferred to liquid nitrogen, cracked open, and then transferred to a fixing solution cooled with liquid nitrogen. Ethanol containing 2% glutaraldehyde and 1% tannic acid was used as a fixing solution. The samples in the fixing solution were kept at −80 °C for 48 h, −20 °C for 2 h, 4 °C for 2 h, and finally returned to room temperature, in that order. Cells on copper disks were immersed in dehydrated ethanol for 30 min thrice and then overnight. Propylene oxide : resin (Agar low viscosity resin LV, cat. no. R1078, Agar Scientific Ltd, Essex, UK) mixture was used for resin replacement of the samples, with the following mixing ratios in sequence; 1 : 0 (30 min, twice), 2 : 1 (1 h, once), 1 : 2 (1 h, once), 0 : 1 (overnight, once). The resin-substituted samples were allowed to solidify for 24 h at 60 °C. Ultrathin sections of 70 nm thickness were then prepared using an Ultracut E ultramicrotome (Reichert-Jung, Austria). The sections were stained with EM Stainer (cat no. 336, Nissin EM, Tokyo, Japan) for 30 min and lead stain (Sigma-Aldrich, MO, USA) for 30 min. The samples were observed using a transmission electron microscope JEM-2100 (JEOL, Tokyo, Japan) equipped with a thermionic electron emission electron gun (LaB6 filament) at an acceleration voltage of 100 kV.

## Abbreviations

MPa: megapascals
CWI: pathway cell wall integrity pathway

## Acknowledgments

We would like to thank Drs. Satoshi Yoshida, Jiro Toshima, Makoto Nagano, and Takuma Kishimoto for providing plasmids; Drs. David Levin and Hidetoshi Iida for providing yeast strains; Dr. Mitsuaki Tabuchi for providing the rabbit polyclonal antibody for Ypk1 and for his critical reading of this manuscript; Mrs. Katsuyuki Uematsu and Yosuke Ishihara for technical assistances on TEM sample preparations; and the laboratory members for facilitating valuable discussions. TEM observation was supported by Center for Instrumental Analysis, College of Science and Engineering, Aoyama Gakuin University.

## Author contributions

Takahiro Mochizuki: Conceptualization and investigation; Toshiki Tanigawa: Conceptualization and investigation; Seiya Shindo: Investigation; Momoka Suematsu: Investigation; Yuki Oguchi: Investigation; Tetsuo Mioka: Cooperation; Yusuke Kato: Cooperation; Mina Fujiyama: Investigation; Eri Hatano: Investigation; Masashi Yamaguchi: Cooperation; Hiroji Chibana: Cooperation; Fumiyoshi Abe: Conceptualization, supervision, writing – original draft, and review and editing.

---

**Fig. S1.**
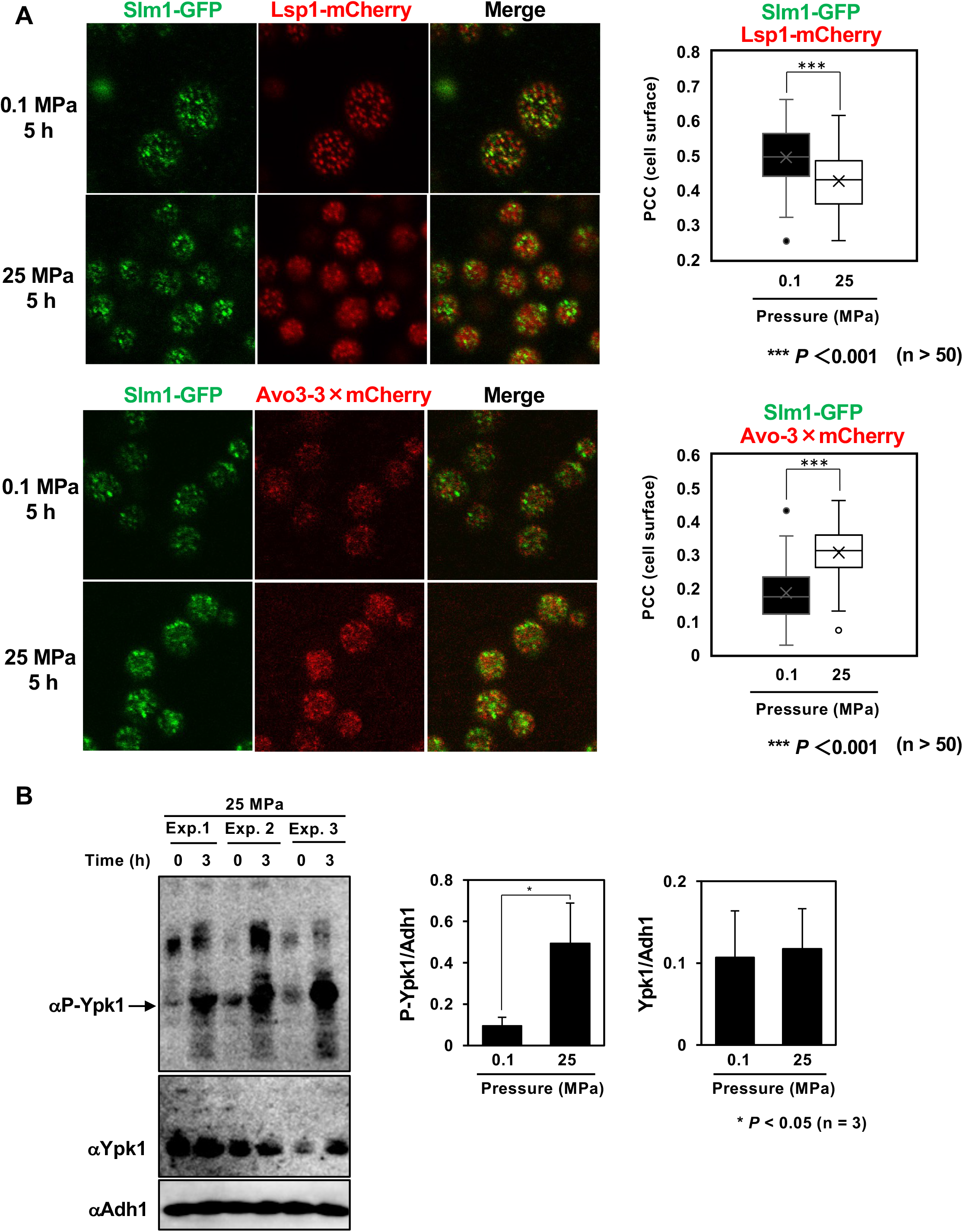
TORC2-mediated Ypk1 phosphorylation by high pressure. (A) High pressure causes the dissociation of a fraction of Slm1 from the eisosomes and its migration to TORC2. Wild-type cells co-expressing Slm1-GFP and Lsp1-mCherry, or those co-expressing Slm1-GFP and Avo3-3×mCherry, were cultured in SC medium at 0.1 or 25 MPa for 5 h. The cells were observed using confocal laser microscopy immediately after depressurization. Colocalizations were analyzed in more than 50 cells from three independent experiments using the cellSens software and represented by the PCC. (B) High pressure promotes the phosphorylation level of Ypk1. Ypk1 and phosphorylated Ypk1 were detected from whole cell extracts via western blotting, using antibodies for Ypk1 and phospho-PKC (pan) (zeta Thr410) (190D10), respectively. The relative intensities, P-Ypk1/Adh1 and Ypk1/Adh1, were quantified in an ImageQuant LAS4000 mini.

